# Phenotypic Differences in Adult and Fetal Dermal Fibroblast Responses to Mechanical Tension

**DOI:** 10.1101/2020.09.30.319749

**Authors:** Walker D. Short, Umang M. Parikh, Natalie Templeman, Oluyinka O. Olutoye, Alexander Blum, Daniel Colchado, Benjamin Padon, Aditya Kaul, Hui Li, Hima V. Vangapandu, Olivia S. Jung, Pranav Bommekal, Phillip Kogan, Monica M. Fahrenholtz, Cristian Coarfa, Swathi Balaji

## Abstract

**Objective:** Fetal regenerative wound healing is characterized by hyaluronan(HA)-rich microenvironment and fibroblasts that produce pericellular matrices(PCM) abundant in high molecular weight HA. Recent studies showed that while small wounds in fetal skin heal regeneratively, large wounds heal with fibrosis. We posit large wounds generate higher mechanical tension which alters HA metabolism in the fetal fibroblasts and lead to a pro-fibrotic phenotype.

**Approach:** C57BL/6J murine fetal (FFB; E14.5) and adult (AFB; 8wk) dermal fibroblasts were subjected to +/-10% tonic strain. Changes in PCM, HA enzymes and molecular weight, and fibrotic gene expression were measured.

**Results:** FFB pericellular matrix reduced upon exposure to increased tension, and the HA profile shifted from high to lower molecular weight. Under static conditions, AFB had higher expression of HA synthases (HAS) 1 and 2 and degradation enzymes KIAA1199, HYAL1, and TMEM2 than FFB, suggesting more HA turnover in AFB. Tension resulted in an increase in HAS1, HAS3, KIAA1199, and HYAL2 expression and a decrease in HAS2 and TMEM2 expression in FFB. CD26, a marker associated with scar production, increased in FFB under tension, along with altered fibrotic gene expression profile and reorganized cytoskeletal f-actin and increased α-SMA that resembled AFB.

**Innovation:** This study elucidates the differences in how biomechanical tension alters HA metabolism and fibrotic phenotype of FFB vs AFB, providing further understanding of the fetal regenerative wound healing phenotype.

**Conclusion:** Understanding the intrinsic differences in HA metabolism and fibrotic phenotype among FFB and AFB in response to wound mechanical stimuli may yield new insights to promote regenerative wound healing.

## INTRODUCTION

Cutaneous wounds in the mid-gestation mammalian fetus heal regeneratively, with wound repair indistinguishable from the surrounding uninjured skin. Significant work has been done to identify the key contributors to the regenerative wound healing responses in the fetus. Among these critical factors are abundant and sustained levels of hyaluronan (HA) along with muted fibrogenic and inflammatory cell and cytokine responses in the wound microenvironment. In addition, regenerative healing is associated with a unique fetal fibroblast phenotype characterized by increased HA production and faster migration and invasion capabilities ^1^. However, the regenerative capacity was not predicated upon the intrauterine microenvironment. Compared to this benchmark, fetuses begin to permanently shift towards a phenotype more prone to scarring in late gestation, which persists throughout the rest of the lifespan. This coincides with increased expression of Engrailed-1 (EN1) and CD26 in fibroblasts in the late gestation and postnatal dermis ^2^. Rinkevich et al. identified these CD26^+^ fibroblasts as the primary drivers of pro-fibrotic remodeling and scar production during wound healing, including in post parturition wounds. However, not all mid-gestation fetal wounds heal regeneratively: larger wounds of critical defect size made in the mid-gestation fetus heal with the formation of a scar, while smaller wounds heal without scar formation ^3^. Specifically, studies done in fetal sheep demonstrate that 50% of full thickness excisional wounds at a wound size of 4-6mm and 6-10mm will form a scar at 80-90 and 60-70-day gestation, respectively ^4^. Beyond an increase in inflammation in large wounds, the underlying mechanisms that govern fetal regenerative healing and their perturbation in large wounds remains only partially understood.

HA is a key extracellular matrix (ECM) component in the wound healing process. HA alters the availability and activity of various growth factors in a wound and regulates the inflammatory and angiogenic responses to injury ^5–8^. The molecular weight of HA dictates its signaling effects. Low molecular weight (LMW) HA exerts pro-inflammatory effects, while high molecular weight (HMW) HA promotes anti-inflammatory responses ^9–12^. HMW-HA has been shown to exert anti-inflammatory effects via the primary cell-surface receptor CD44. HA expression in the wounds is governed by synthesis enzymes hyaluronan synthase (HAS) 1-3 and degradation enzymes hyaluronidase (HYAL) 1-2, KIAA1199, and TMEM2 ^13, 14^. Importantly, HAS1-2 are associated with HMW-HA production (2×10^5^ – 2×10^6^ kDa), while HAS3 is associated with the production of relatively lower molecular weight (1×10^5^ – 1×10^6^ kDa) HA ^15^. HMW-HA stability is critical to sustain and promote its role in regenerative wound healing, as its catabolism in the wounds generates LMW-HA fragments that promote pro-inflammatory and pro-fibrotic responses ^16^. Thus, the balance of HA-forming synthases and HA-degrading enzymes impacts the final balance of HMW-to LMW-HA in the wound microenvironment, which may consequentially affect how the wound heals.

Dermal fibroblasts are the functional cells of wound repair that drive both the synthesis and remodeling of the wound ECM ^1^. While different fibroblast lineages are currently being delineated, the phenotype and activity of mid-gestational fetal dermal fibroblasts (FFB) differ significantly from the phenotype of late gestation and postnatal/adult dermal fibroblasts (AFB) ^2, 17^. Notably, these phenotypic differences include their migratory potentials, rate of proliferation and differentiation, cytokine production, and ECM synthesis ^18^. More specifically, we have shown that FFB secrete more HA than AFB and have large HA-rich pericellular matrix coats (PCM) around them ^19^. This pericellular coat of HA both regulates the metabolic rate of FFB and facilitates their migration and invasion. Importantly, inhibiting HA production in FFB using 4-methylumbeliferone (4-MU), an HA synthase inhibitor, reduced their PCM and attenuated their levels of migration and invasion to levels similar to that of AFB ^19^. These differences in fibroblast HA regulation may partially explain why mid-gestation mammalian fetuses heal regeneratively, whereas similar wounds in the post-natal period produce fibrosis and heal with scar.

Research demonstrates that wound tension alters the expression of fibrogenic markers in fibroblasts. Fibroblasts in a wound are subjected to changes in mechanical force balance during wound healing, and respond by generating traction forces to close the wound by bringing the wound margins together ^20, 21^. Mechanically stressed fibroblasts differentiate into a specialized myofibroblast phenotype that is characterized by expression of α-smooth muscle actin, formation of prominent stress fibers, and collagen production leading to fibrosis and contraction ^22, 23^. In practice, this means tension on a wound should be minimized to lower the risk of scarring. Indeed, offloading tension on adult skin wounds significantly reduce scarring, relative to wounds with no stress shielding ^24^. Previous studies have shown that the resting tension in fetal skin is negligible and significantly lower than that in adult skin ^25^. The lower resting tension of fetal skin may act much like a stress-shield and is perhaps linked to regenerative healing of small and moderate sized wounds. However, increasing wound defect size in the fetus shifts the wound healing process from regenerative to a more inflammatory and scarring phenotype through unknown mechanisms. We posit that increasing wound size in fetal wounds increases the mechanical forces on the fibroblasts and the tensile forces needed to close the wound. However, the fetal fibroblast responses to increases in mechanical tension are not known.

In sum, evidence shows that the balance of HMW-HA and LMW-HA within a wound and the mechanical tension in a wound can individually determine the fibrotic outcome. However, it is currently not well understood how increases in mechanical loads during wound healing alter FFB HA metabolism. Thus, we sought to determine how mechanical tension influences the balance of HA turnover and the fibrogenic phenotype of fetal fibroblasts as compared to AFB. We hypothesize that increasing mechanical tension on FFB alters HA metabolism, with tension driving production of LMW-HA and profibrotic responses in FFB.

## CLINICAL PROBLEM ADDRESSED

Dysregulated healing, including scarring and impaired healing in chronic wounds, results in a significant socioeconomic and psychosocial burden. Contrarily, regenerative healing culminates in tissue with restored integrity, function, and cosmesis. Research leading to true regenerative healing is of paramount importance to clinicians who care for any patient with a wound. Fetal dermal wound healing is the paradigm of this regenerative phenotype, and uncovering the biologic underpinnings of this phenotype will advance development of wound healing therapeutics. Notably, our data support the notion that fetal fibroblasts shift their metabolism of hyaluronan in response to tension, which also alters their fibrotic gene expression profile towards that of adult fibroblasts.

This data further advances our understanding of the behavior of these regenerative cells in the wound environment.

## MATERIALS AND METHODS

### Cell isolation and culture

Primary fetal murine dermal fibroblasts (FFB) were isolated from mid-gestation age C57BL/6J fetal mice (E14.5, Strain number 000664, Jackson Laboratories, Bar Harbor, ME) and adult murine dermal fibroblasts (AFB) were isolated from C57BL/6J adult mice (8–12 weeks) using standard isolation protocols as previously described ^26^. Dorsal skin was excised and used for cell isolation. Fibroblast cultures were maintained in Dulbecco’s Modified Eagle’s Medium with GlutaMAX 1mg/mL DMEM (GIBCO, Carlsbad, CA) supplemented with 10% bovine growth serum (BGS) (Hyclone, Logan, UT), 1% sodium pyruvate, 1% penicillin (10,000 U/mL stock), and 1% streptomycin (10,000 μg/mL stock) (Invitrogen, Carlsbad, CA). Cultures were maintained in a humidified chamber at 37 °C with 5% CO2. Cells between passages 3–5 were used for experiments; upon thawing, cells were acclimated to either plastic or silicone membrane for 1 passage pior to experiments. Cells between groups were equal in passage for individual experiments, n=3 experimental replicates, each from a passage between 3-5. All protocols were approved by the Baylor College of Medicine Institutional Animal Care and Use Committee.

### Fibroblast cultures on flexible membranes

A mechanical strain device (FX3000, Flexcell International Corp.) was used to apply cellular strain to murine FFB and AFB to determine the effect of mechanical tension on their HA production and fibrotic phenotype. Cells were seeded as monolayers on silicone membranes in 6-well plates (BioFlex membrane Culture Plate, Flexcell Corp.). Per the manufacturer’s material properties specification, the Young’s modulus of the silicone membranes is 930 kPa, which provides a softer substrate for cell growth when compared to standard plastic culture dishes that have a Young’s modulus of 1×10^7^ KPa and better mimics in situ conditions. Prior to cell seeding, and to improve cell adherence to the membrane, the membranes were incubated for 24 hours with 1% gelatin, which was then aspirated. While this surface coating allows better cell attachment, it does not affect the material properties of the membrane. 75,000 cells in 3 mL of fresh Dulbecco’s Modified Eagle’s Medium containing 10% BGS were added to each well, and cells were allowed to attach to membranes overnight. The next day, culture medium was exchanged, 3 mL of fresh Dulbecco’s Modified Eagle’s Medium containing 2% BGS was added, and cells were cultured in the reduced serum conditions for 24 h. After this period, tension was initiated. Six planar cylinders (25 mm diameter) served as loading posts for each of the 6 wells in the culture plates, which were centered beneath each 35-mm well. By application of vacuum, equibiaxial strain at 10% was applied to the fibroblast monolayers, which underwent tonic deformation for 1, 3, 6, 12 and 24 h. Cells cultured in identical conditions on the flexible membranes without applied tension in the same incubator served as static culture controls.

### Particle Exclusion Assay for Pericellular Matrix Analysis

Pericellular matrix (PCM) area was measured using a particle exclusion assay as described previously ^26^. Briefly, AFB and FFB cultures were rinsed with PBS, 1mL glutaraldehyde-stabilized sheep erythrocytes (1 × 10^8^ cells/mL; Intercell Technologies, Jupiter, FL, USA) were added to each well and allowed to settle for 10 min, and the exclusion of the fixed red blood cells was used to visualize fibroblast pericellular coats. Individual cells were imaged under phase contrast at 20x magnification with an Axiovert 100M inverted phase contrast microscope (Carl Zeiss, Thornwood, NY, USA), which showed the outline of the cell body and the outline of the HA coat as a halo around the cells bounded by the red blood cells. Images of randomly selected cells from each group (20 cells/well) were analyzed with ImageJ software, and the PCM ratio was reported as the ratio of the area of the PCM plus the cell body area to the cell body area alone. Each experiment was conducted with duplicate wells per group, and all experiments were repeated 3 times.

### Immunocytochemistry

After 6 h and 24 h under static or tension conditions, cells on the bioflex membrane plates were fixed with 4% paraformaldehyde for 10 minutes at room temperature. Cells were then permeabilized with 0.01% Triton-X100 when looking for intracellular targets only and blocked with 1% bovine serum albumin (BSA) in PBS for 1 h at room temperature. Primary antibodies were diluted in 1% BSA and allowed to bind for 1 h at room temperature. For detection of fibrotic markers, cells were incubated with anti-CD26, (rabbit mAb ab28340; Abcam, Cambridge, MA) or anti-α-SMA (rabbit mAB ab5694; Abcam), then detected with an appropriate fluorescent secondary antibody for 1 h at room temperature. For detection of HA, cells were similarly incubated with HA binding protein (HABP) (EMD Millipore 385911, Burlington, MA), then detected with an appropriate fluorescent secondary antibody for 1 h at room temperature. Phalloidin (ab176753, Alexa Fluor 488 conjugated; Abcam, Cambridge, MA) or Wheat Germ Agglutinin (ab178444, Alexa Fluor 488 conjugated, ThermoFisher) were used to detect cell filamentous actin and cell membrane respectively, and samples were counterstained with DAPI. After completion of the staining, the bottom surface of the membranes that were in contact with the loading posts were first gently cleaned with a cotton tip applicator to remove the residual grease that is coated on this side of membranes to reduce friction during the tension cycles (as per the manufacturer’s protocol). The membranes were then cut out and mounted on a glass slide for imaging with a DMi8 inverted fluorescent microscope (Leica Microsystems, Buffalo Grove, IL).

### HA Extraction and Gel electrophoresis

HA extraction and gel electrophoresis were performed, as previously described ^27^. Briefly, cell culture supernatants were collected after treatment, and 10% volume of Proteinase K (Invitrogen, 1 mg/mL with SDS at 0.01%) was added and incubated at 60 °C for 4 hours to digest the proteins. 4 mL pre-chilled (-20 °C) 200 proof ethanol was added and incubated overnight at −20 °C. Samples were then centrifuged at 14,000g for 10 minutes at room temperature (RT), and the pellets were washed with 4 mL 75% EtOH, vortexed, and centrifuged at 14,000g for 10 minutes. The supernatant was discarded, and the pellet was air dried at RT for 20 minutes. 100 μL of 100 mM ammonium acetate (Sigma-Aldrich) was added to re-suspend the pellet, and the samples were heated at 100 °C for 5 minutes to inactivate Proteinase K. Samples were then chilled on ice for 5 minutes, followed by addition of 1 μL of Benzonase (EMD Millipore) and incubation overnight at 37 °C to digest nucleic acids. Samples were boiled for 5 minutes to inactivate enzymes. We then added 200 μL of cold 200 proof ethanol to the suspension and incubated overnight at −20 °C, followed by centrifugation at 14,000g for 10 minutes at RT. Supernatant was discarded, and the pellet was washed with 1 mL cold 75% ethanol, centrifuged as before, and air dried for 20 minutes. The pellet was then re-suspended in 10 μL of 10 M formamide and kept at 4 °C overnight prior to electrophoresis.

A 1% agarose gel was used for HA gel electrophoresis (SeaKem HGT Agarose, Cambrex). 2 μL of the sample loading solution (0.2% bromophenol blue in 10 M formamide) was mixed with each 10 μL extracted HA sample. Hyaluronan standards (Select-HA^TM^, Hyalose) were prepared similarly. Prior to loading, HA concentration was confirmed in the extracted samples with a Hyaluronic Acid test kit (Corgenix Specialists in ELISA & IVD Diagnostic Technology) according to the manufacturer’s instructions to load equal amounts. Samples and standards were loaded, and the gel was run at 100 V until the tracking dye migrated about 75% of the length of the gel. Gels were equilibrated with 30% ethanol, then stained with Stains-All (6.25 μg/mL) (Sigma-Aldrich) solution overnight in the dark. Gels were washed and equilibrated with water before de-staining via light exposure followed by imaging.

### Real-time Quantitative PCR and PCR Fibrosis Array

After completion of the mechanical loading cycle for 1, 3, 6, 12 or 24 h, culture supernatants were collected and stored at −80 °C with protease inhibitor cocktail (Sigma). Cell monolayers were washed twice in PBS. PBS was aspirated, and RNA lysis buffer was added to the wells in both tension and static control cultures. A cell scraper was then used to dissociate cells, only RNA obtained from adherent cells was used, free-floating cells were washed away and not analyzed. Lysates were vortexed and frozen at −80 °C. Total RNA was isolated using the PureLink^TM^ RNA Micro Kit (Invitrogen). After concentration and purity determination with a nanodrop, cDNA was reverse-transcribed from 500 ng of RNA using the high capacity RNA to cDNA kit (Cat # 4387406) following the manufacturer’s protocols (Applied Biosystems, Foster City, CA, USA) and amplified by quantitative real time PCR using forward and reverse primers (Integrated DNA Technologies, Coralville, IA, USA) and Power SYBR Green PCR Master Mix in a Bio-Rad CFX 384 real-time system. Mouse RPS20 gene expression was used to normalize the amount of the amplified products. Relative expression values were calculated by using the ΔΔCt method ^28^. Primer sequences are listed in **Table 1**; those purchased from Bio Rad are indicated.

**Table 1.**
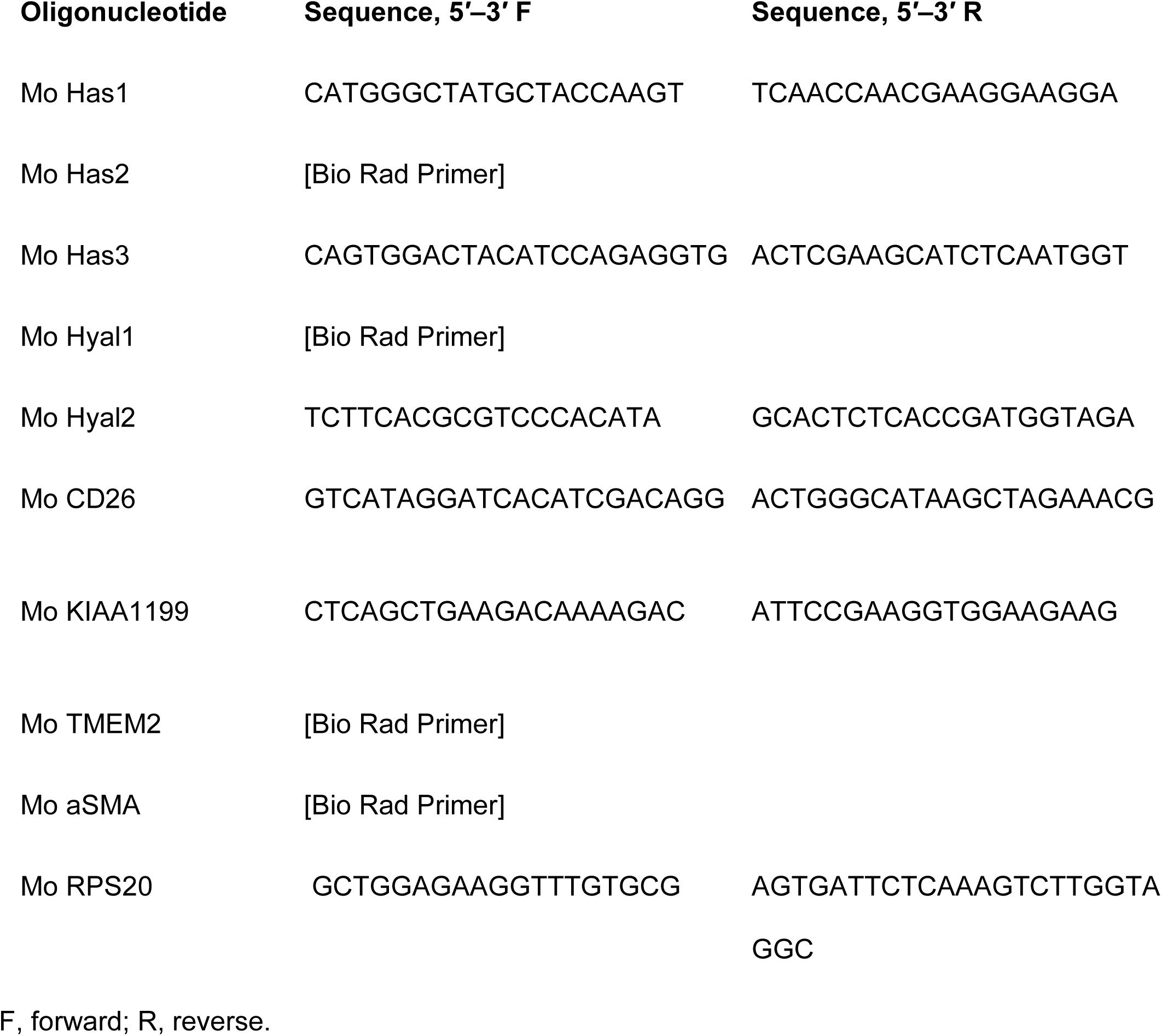
Primer sequences for quantitative real time PCR validation.

We also used the RT^2^ Profiler PCR array mouse fibrosis (cat. #PAMM 120-ZE-4) kit from Qiagen to detect the expression of 84 fibrosis-related genes. This assay uses cDNA made with RT^2^ first strand kit (Qiagen cat. #330404) and RT^2^ SYBR Green qPCR Mastermix (Qiagen cat. #330501). Bio-Rad CFX 384 real-time system and Bio-Rad CFX manager software were used for real-time PCR. Data were analyzed by Qiagen RT^2^ Profiler PCR Array data analysis software. Principal Component Analysis (PCA) and multi-dimensional sampling were performed with Orange software (https://orange.biolab.sion) on the ΔCT values for the genes. Venn diagram in Figure 6C was created using any up-or down-regulated gene, while Tables 3 and 4 are composed only of genes in which the fold changes was greater than 2 in the positive or negative direction.

**Table 2.** Overall changes in regulation of HA synthesis and degradation genes in adult (AFB) and fetal (FFB) fibroblasts after tension. A combination of these gene expression changes result in an overall increase in lower molecular weight HA.

|  | **AFB +**<br>**Tension** | **FFB +**<br>**Tension** |
| --- | --- | --- |
| HAS1 (HMW) | ↑ | ↑ |
| HAS2 (HMW) | ↓ | ↓ |
| HAS3 (LMW) | ↑ | ↑↑ |
| HYAL1 | ↓ | ↓ |
| HYAL2 | --- | --- |
| KIAA1199 | ↑ | ↑↑ |
| TMEM2 | ↓ | ↓ |
| **Cumulative**<br>**Effect** | **LMW HA** | **LMW HA** |

**Table 3.** List of genes that are commonly and uniquely upregulated in adult (AFB) and fetal (FFB) fibroblasts under tension as compared to static culture conditions.

| Upregulated genes |  | Fold Regulation |  |
| --- | --- | --- | --- |
| Gene Symbol | Gene Name | AFB | FFB |
| *Common* |  |  |  |
| FasI | Fas Ligand | 14.26 | 7.45 |
| IL5 | Interleukin 5 | 9.74 | 2.87 |
| IL13 | Interleukin 13 | 3.22 | 4.61 |
| IL10 | Interleukin 10 | 2.88 | 32.46 |
| Jun | c-Jun | 2.81 | 2.01 |
| IL1a | Interleukin 1a | 2.34 | 2.19 |
| *Only in AFB* |  |  |  |
|  | TIMP metalloproteinase |  |  |
| TIMP4 | inhibitor 4 | 8.09 |  |
| Agt | angiotensinogen | 5.57 |  |
| *Only in FFB* |  |  |  |
| MMP13 | matrix metalloproteinase 3 |  | 8.95 |
| IL4 | Interleukin 4 |  | 6.01 |
| Cxcr4 | CXC motif chemokine receptor 4 |  | 4.67 |
| Bmp7 | bone morphogenetic protein 7 |  | 2.56 |

**Table 4.**
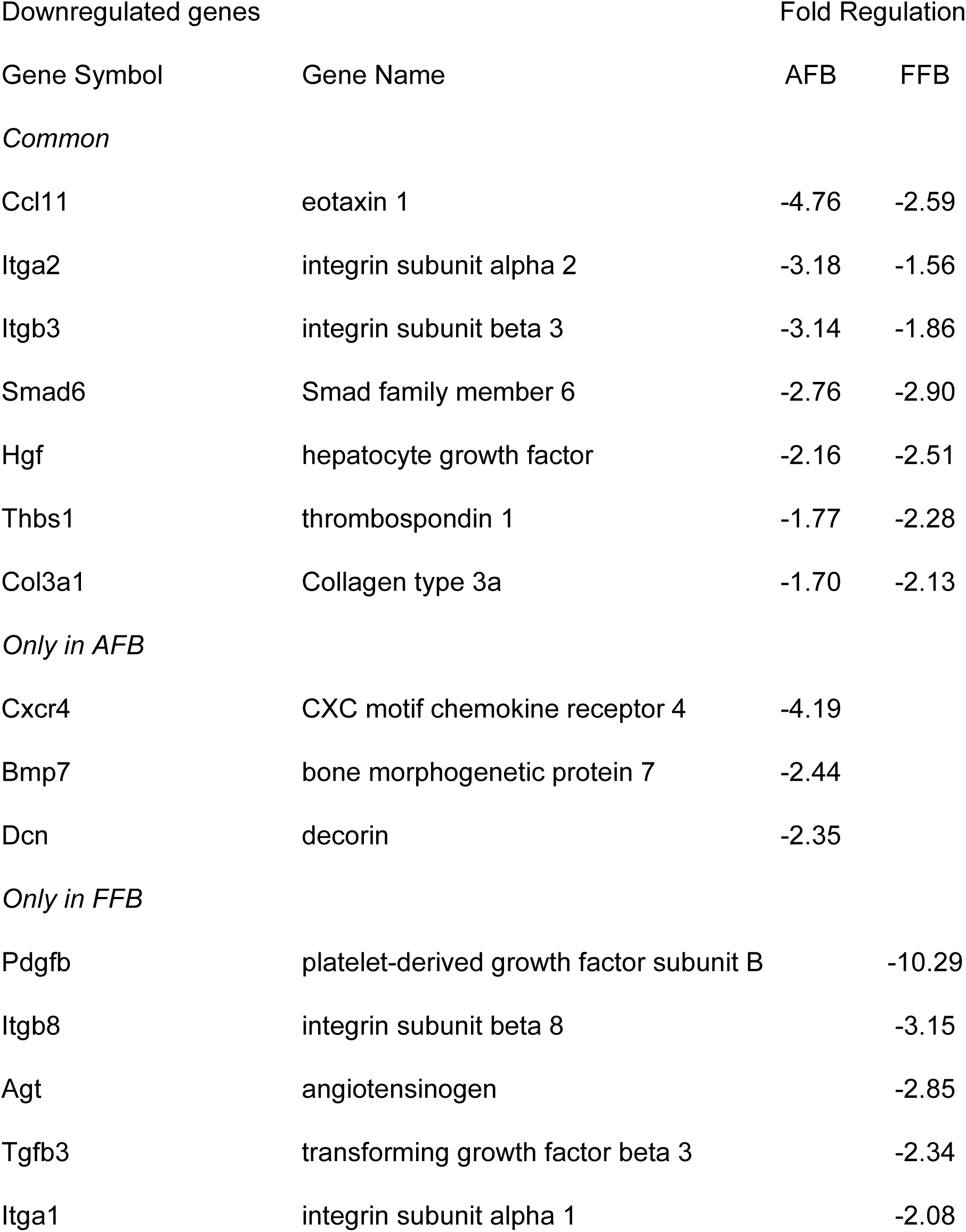
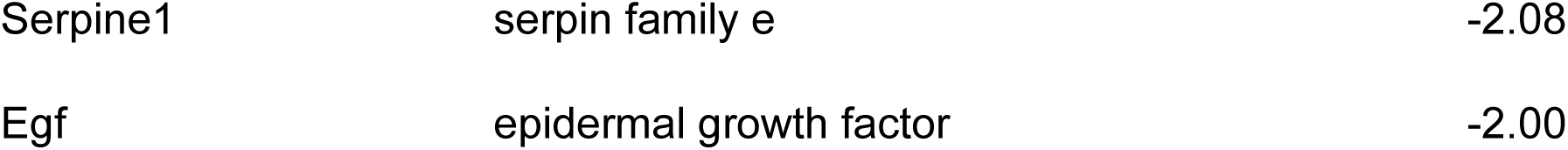
List of genes that are commonly and uniquely downregulated in adult (AFB) and fetal (FFB) fibroblasts under tension as compared to static culture conditions.

### Protein extraction and Immunoblotting by ProteinSimple

AFB and FFB from static and tension conditions were washed twice with PBS and lysed in RIPA buffer (Thermo Fisher Scientific, Waltham, MA, USA), and digested samples were subjected to ProteinSimple Wes System (Biotechne, San Jose, California)^29^ with a 12-230 kDa separation module (ProteinSimple SM-W004), a new automated capillary-based quantitative electrophoresis system that is reliable, reproducible, and accurate where the signals are automatically recorded, quantified, analyzed and visualized using Compass software. Protein was only obtained from adherent cells, free-floating cells were washed away and not analyzed. Primary antibodies are anti-SMA (Abcam5694, Cambridge MA, USA) and anti-Beta Actin (Cell Signaling 3700, Danvers MA, USA) and diluted as 1: 500 and 1: 200, respectively. Secondary antibodies are ProteinSimple Anti-Rabbit Streptavidin-HRP (ProteinSimple DM-001), and Anti-Mouse Streptavidin-HRP (ProteinSimple DM-002, Biotechne, San Jose CA, USA). The assay was run according to the manufacturer’s directions with no deviations, and protein bands were visualized and analyzed with ProteinSimple Biotin Detection Module (ProteinSimple DM-004). The sample analyte concentrations were derived through calculating the ratio of area under the peak for anti-SMA and beta actin in each sample.

### Imaging and Quantification of Cell Numbers

In order to identify whether there was a change in cell density after tension, we quantified the number of attached cells after static and tension treatments in both AFB and FFB. Cells were plated as described above on 6-well flexible membrane plates. Cells were then subjected to either static conditions or 3, 6, 12 or 24 hours of 10% tonic strain. After completion of the loading cycle at each time point, the wells were washed with PBS twice and the number of cells still adherent to the membrane were counted. Five phase contrast images were taken (Leica DM IL LED) per well from the different groups. Images were quantified using Fiji Software ^30^, and only cells that maintained spindle-like structure were included in the counts. Counts were obtained from duplicate wells per group for each timepoint, and the experiments were repeated twice.

### Detection of Cell Viability with Lactate Dehydrogenase (LDH) Assay and Cell Counting with Trypan Blue

In order to quantify cell death after tension, we utilized the LDH assay to detect levels of lactate dehydrogenase, an enzyme that is present in the cytosol of cells and is released into culture media, when the integrity of the cell membrane is lost or disrupted leading to cell death. Similar to above, cells were seeded in a 6-well flexible membrane plate at a density of 75,000 cells per well in 3 mL DMEM containing 10% BGS and allowed to adhere overnight. The media was removed, and cells were serum starved in DMEM containing only 2% BGS for 24 hours prior to treatment. Cells were then subjected to 6 hours of 10% tonic mechanical strain. Cells were cultured in identical conditions on the flexible membranes without applied tension in the same incubator served as static controls. Immediately prior to and after tension application, 200 µL of cell culture supernatant was collected and the Cytotoxicity Detection Kit (LDH) (Roche, Indianapolis, IN) was then utilized to detect LDH levels in the culture supernatant. The remaining supernatant was used to detect the viability of cells floating in the supernatant using trypan blue staining.

Briefly, for the LDH assay, reagents were prepared according to manufacturer’s instructions, and 100 µL of culture supernatant was combined with 100 µL of assay reagent and incubated for 30 minutes. The absorbance was then measured at 490 nm. For both AFB and FFB, all absorbance readings were normalized to that of DMEM media with 2% fetal bovine serum (FBS). Final values were expressed as the absorbance reading from tension conditions normalized to static control conditions for each cell line at each time point. For cell counting with trypan blue, 50 µL of cell culture supernatant was combined with 50 µL of Trypan Blue (0.4% Solution, Thermo Fisher Scientific Inc, Waltham, MA), and 10 µL was used for cell counting using a hemocytometer. Live cells were defined as cells that excluded the stain, and dead cells were defined as cells that positively took up the stain. Similar counting of the floating cells in the cell culture supernatant of static control plates was performed. Each experiment was conducted with duplicate wells per each group, and the experiments were repeated twice.

### Live/Dead Cell Staining

We further utilized the fluorescent Live/Dead Fixable Near-IR Dead Cell Stain Kit (Invitrogen, Carlsbad, CA) to determine the viability of the cell monolayers after tension. After 6 hours of 10% tonic tension, cells on flexible membranes from the tension treated well and static control wells were washed with PBS. 1 mL of DMEM media with no FBS was added to each well, along with 1 µL of fluorescent reactive dye (prepared according to manufacturer’s instructions). Cells were incubated for 30 minutes, washed with PBS twice, and then fixed with 37% formaldehyde for 15 minutes. Cells were again washed with PBS three times. Membranes with fixed cells were cut and mounted on charged slides. Slides were then imaged with Keyence BZ-X800 microscope (Keyence, Osaka, Japan) using Brightfield and Texas Red filters. Viable cells have less intense fluorescent detection, as the dye’s reactivity is restricted to the cell-surface amines, resulting in less intense fluorescence. As a positive control, cell monolayers on the flexible membrane were subjected to intense mechanical swabbing of the membrane on the non-cell coated side resulting in cell membrane compromise, which would allow the dye to react with free amines both in the cell interior and on the cell surface, yielding intense fluorescent staining. Each experiment was conducted with duplicate wells per group, and the experiments were repeated twice.

### Statistical Analysis

Statistical significance of the data was evaluated using analysis of variance (ANOVA), followed by post-hoc tests (Tukey) and Student’s t-test when appropriate in MS-Excel (Microsoft, Redmond, WA).

## RESULTS

### Tension decreases PCM surrounding fetal fibroblasts

We have previously shown that the FFB produce a characteristic HA-rich PCM coat on the cell surface, and when HA was inhibited with addition of either 4-MU or hyaluronidase to FFB cultures, their PCM coats were reduced and resembled those in AFB, both in the size of the coat and their SEM structure ^18, 26^. We therefore used PCM analysis to interrogate the impact of tension on extracellular matrix production by FFB, illustrated in **Figure 1A**. Baseline PCM measurements demonstrate significantly decreased PCM surrounding AFB when compared to FFB (1.84 ± 0.08 PCM area/Cell area vs 2.78 ±0.14 PCM area/Cell area, p<0.01), which is consistent with prior findings. After 24 hours of tension, FFB cultured under tension showed a reduced PCM area as compared to FFB under static conditions (1.98 ± 0.15 PCM area/Cell area vs 2.78 ± 0.14 PCM area/Cell area, p<0.01) and resembled the PCM of AFB grown in static conditions (1.98 ± 0.15 PCM area/Cell area vs 1.84 ± 0.08 PCM area/Cell area), while AFB showed a small increase in PCM under tension **(Figure 1B).**

**Figure 1:**
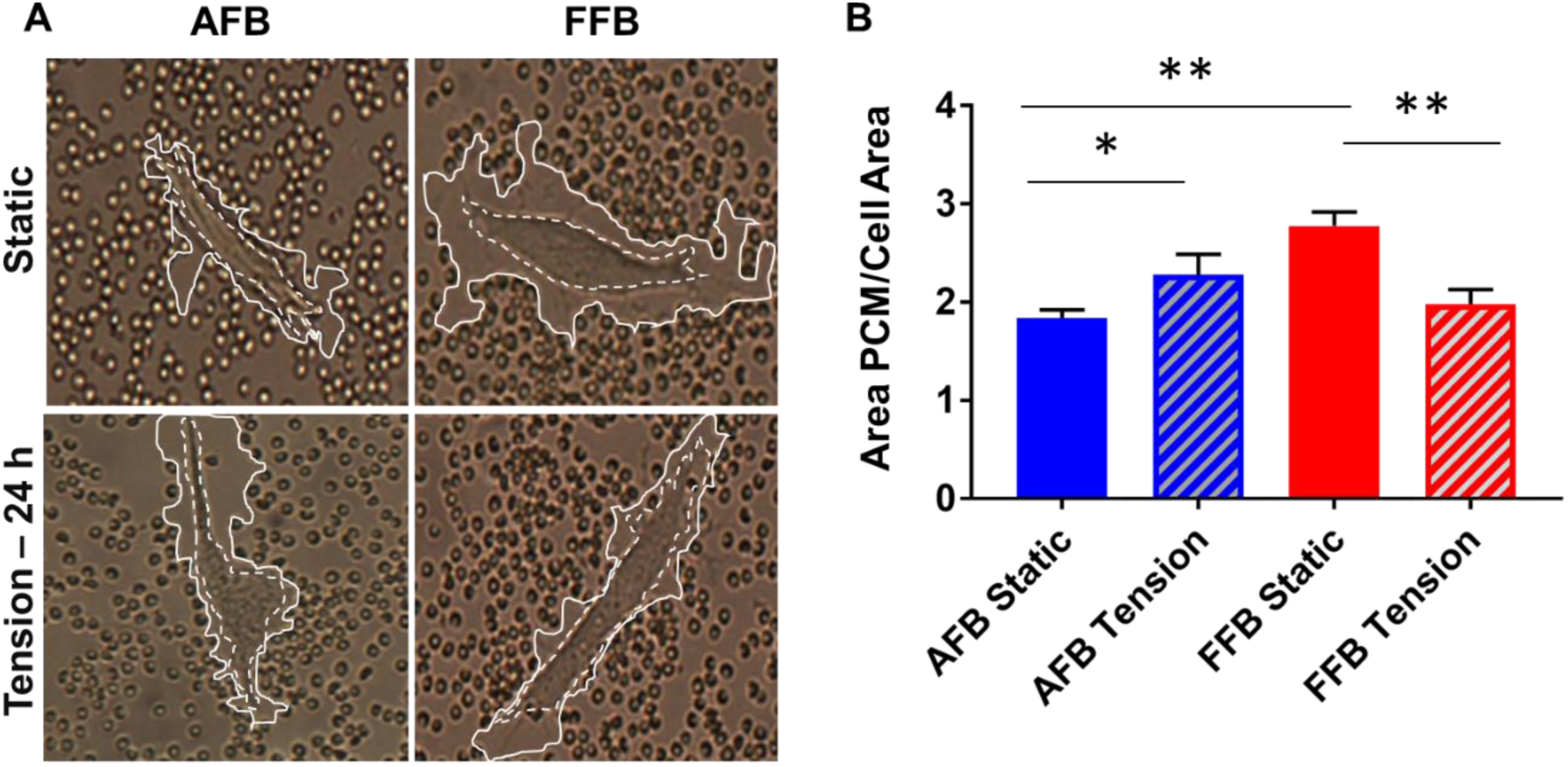
(A) Representative images of the pericellular matrix coat (white outline) secreted by adult (AFB) and fetal fibroblast (FFB) under static and 24 h tension conditions. Dotted white line trace the cell body. (B) Quantification of PCM area over cell area. n = 4-6 experimental replicates. *p<=0.05, **p<=0.01. Graphs are reported as mean ± SD.

We further explored changes in HA expression in FFB exposed to tension by staining these cells for HA. Representative staining for HABP with appropriate controls **(Supplemental Figure 1A-D)** in FFB after 6 hours of tension compared to static culture are shown. In static control FFB, HABP staining was robust and appeared to have rope-like protrusions from the cell surface that are connecting the different cells, as shown by the arrows in **Supplemental Figure 1A.** After 6 hours of tension, these rope-like protrusions disappeared and long thin tendrils between cells were seen as shown in **Supplemental Figure 1B.**

### Tension shifts HA metabolism toward LMW-HA

Knowing that the majority of the PCM surrounding the FFB consists of HA, which we previously demonstrated with gain and loss of function experiments ^18^, we sought to understand how tension impacts HA metabolism to further explain the observation that tension reduces the PCM coats of FFB. Expression profiles of HA metabolism-related genes (HAS1-3, KIAA1199, HYAL1-2, and TMEM2) at intervals of 0, 1, 3, 6, and 12 hours after the application of tension were measured by qRT-PCR **(Figure 2).** FFB and AFB demonstrate different HA metabolism profiles at baseline 0 h, with both significantly higher synthase HAS1 (1.004 vs 6.8 RQ, p<0.05), HAS2 (1.003 vs 1.7 RQ, p<0.05), and hyaluronidase KIAA1199 (1.006 vs 17.5 RQ, p<0.05), HYAL 1 (1.02 vs 4.4 RQ, p<0.05), HYAL 2 (1.003 vs 1.4 RQ, p<0.05), and TMEM2 (0.966 vs 1.398, p<0.01) expression in AFB when compared to FFB **(Figure 2A).** This suggests that, at baseline, AFB may have higher PCM turnover.

**Figure 2:**
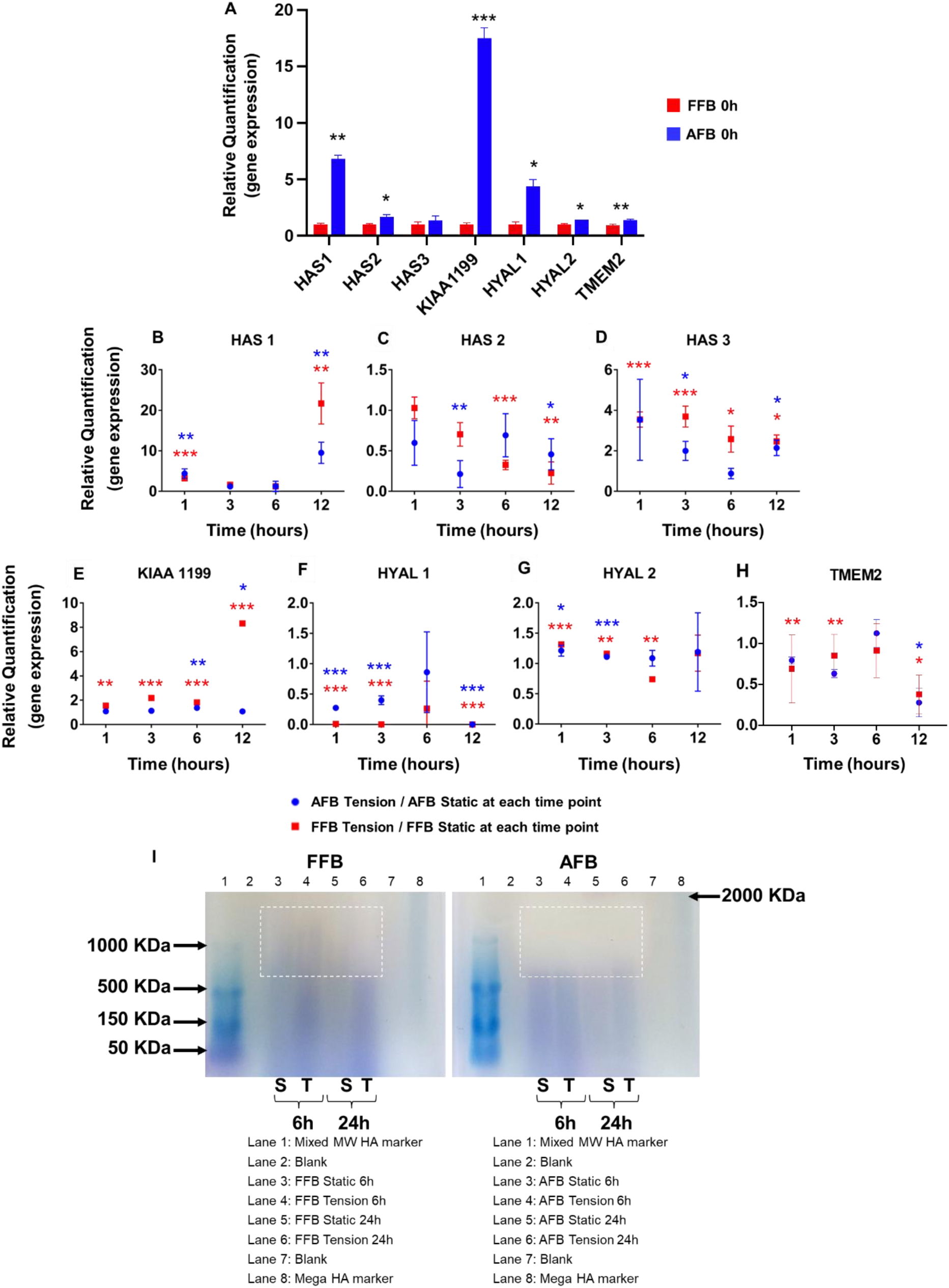
Regulation of HA metabolism over the course of 24 h of static and 10% tonic tension conditions. (A) Baseline differences in the gene expression of the main HA synthases (HAS1-3) and hyaluronidases (HYAL1-2, KIAA1199, and TMEM2) between AFB and FFB prior to the application of tension are shown. Data is reported as mean ± SD, *p<0.05. (B-D) Effect of tension on the relative expression of hyaluronan synthase 1-3, (E) KIAA1199, (F-G) hyaluronidases 1 and 2, and (H) TMEM2 are shown. At each time point, 1, 3, 6, and 12 hours, the relative quantification of each of the genes of interest in tension conditions as compared to the paired static controls for each cell type are expressed as mean ± SD. *p<=0.05, **p<=0.01, ***p<=0.001 denotes significant changes in tension compared to baseline expression in each cell line. Blue * indicates significance value for AFB, Red * indicates significance value for FFB. N = 3 experimental replicates. (I) Electrophoresis gel indicating total hyaluronan molecular weight profiles in either FFB (left) or AFB (right) cell culture supernatant in static and tension conditions at 6 and 24 hours. AFB, regardless of static or tension conditions, shown no HMW-HA smear above 900KDa. In contrast, FFB under static culture conditions or with short period (6 h) of tension express HMW-HA, but have negligible expression of HMW-HA smear (indicated by the dotted line box) after 24 h tension and are similar in molecular weight expression profile to AFB.

We then determined the relative increased/decreased expression of each of the genes of interest in tension conditions compared to their static controls at each time point 1, 3, 6, and 12 hours for both FFB and AFB to evaluate how each cell type responds to tension over time. Tension had differential effects on HA-related gene expression in FFB and AFB **(Figure 2B-G).** Specifically, HAS1 expression increased with tension compared to static conditions, and while there was no difference in this increase between AFB vs. FFB at 1, 3, or 6 h, we noticed that at 12 h FFB had 20-fold increase over the FFB static control, while AFB showed only a 10-fold increase over its static control. HAS3 expression also increased in both FFB and AFB with tension. While there was a decline noted over time in AFB, FFB maintained increased expression under tension as compared to its static at 3 and 6 h. KIAA1199 expression increased with tension in FFB, with almost an 8-fold increase at 12 h, while remaining unchanged in AFB. By contrast, there was decreased HAS2 expression in both AFB and FFB with tension, with the effect of tension in AFB more pronounced than in FFB at 3 h as compared to their respective controls. HYAL1 and TMEM2 expression decreased over time in both FFB and AFB under tension, whereas HYAL2 expression slightly increased in AFB and FFB at 1 and 3h, and it remained increased over time in AFB but showed a slight decrease in FFB at 6 h. In sum, tension resulted in an overall increase in HAS1, HAS3, KIAA1199 and HYAL2 expression, and decreased HAS2 and HYAL1 for FFB and AFB under tension, albeit at different levels **(Table 2).**

Keeping in view the changes in both the HA synthases and degradation enzymes, we sought to determine the effect of the tension on HA molecular weight patterns using gel electrophoresis. As opposed to specific molecular weight of HA used for the ladder that produces distinct bands at each predicted weight, HA in biologic tissues is constantly turning over and comprises a mix of molecular weight variants that appear as a smear. Samples that have HMW-HA produce a smear between the 2000KDa and 1000KDa bands. Images of the gels **(Figure 2I)** demonstrate that the HMW-HA (>9×10^5^ kDa) smear pattern noted in the FFB cultures at 6 hours +/-tension is by-and-large absent after 24 hours of tension in FFB, which resemble the molecular weight of HA produced by static AFB and AFB after both 6 and 24 hours of tension. The breakdown of HMW-HA in FFB in response to 24 hours of tension resulted in an increased staining intensity in the lower molecular weight range between 50-500 KDa.

### Tension leads to reorganization of actin in fetal fibroblasts

As the changes in the PCM coats of the FFB may affect the cell’s interaction with its microenvironment and thereby drive potential changes in its internal cytoskeletal organization, we sought to understand the effect of tension on the organization of actin. AFB and FFB were stained for filamentous actin using fluorophore-conjugated phalloidin after 6 hours of static and tension conditions. Qualitatively at baseline, F-actin in static AFB appears as long filaments running in a parallel orientation, in contrast to the unorganized appearance composed of shorter cords in static FFB (**Figure 3**). Following 6 hours of tension, F-actin in FFB shifts into a more parallel orientation of longer cords, akin to AFB under both static and tension conditions.

**Figure 3:**
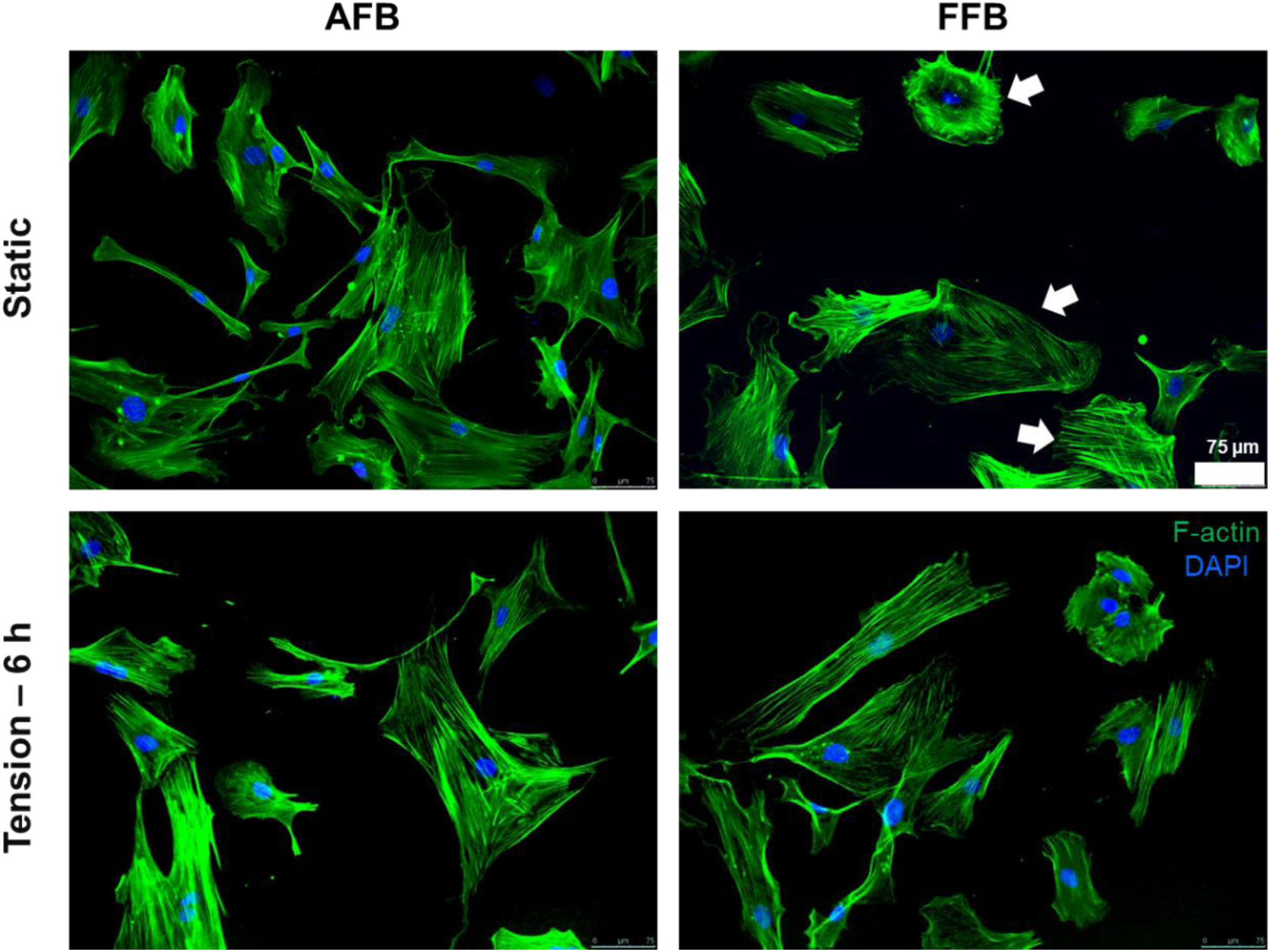
Immunocytochemistry fluorescent staining of F-actin (Phalloidin stain, green fluorescence) in AFB and FFB under static and 6 h tension conditions is shown. Shorter and disorganized F-actin filaments in FFB under static conditions are indicated by arrows, which differ from the parallel fiber orientation in AFB and FFB under tension. Blue stain indicates DAPI staining for cell nucleus. Scale bars represent 75 µm.

Staining for α-SMA was in congruence with these findings. AFB and FFB were stained with antibodies against α-SMA after 24 hours under static and tension conditions to allow for protein expression. At baseline in static culture, we observed more α-SMA expression in AFB than FFB **(Figure 4A).** α-SMA fibers in static AFB appear to be in a parallel orientation, in contrast to the disorganized fibers seen in static FFB. Following 24 hours of tension, we qualitatively observed more α-SMA in AFB, and while we did not note an overall increase in α-SMA expression in FFB, the fibers appeared to be organized in the same parallel fashion as AFB. Expression of α-SMA by AFB was confirmed to be significantly increased at multiple timepoints (1, 3, 6, and 12h) under tension using qRT-PCR analysis, while no significant change in gene expression of α-SMA was seen in FFB when exposed to tension at these timepoints **(Figure 4B).** Protein levels of α-SMA did not significantly change with tension in FFB compared to static control. While levels of α-SMA did increase in AFB following tension, approaching statistical significance (p=0.07), there was a significant difference between α-SMA produced by AFB and FFB under tension (p<0.05) (**Figure 4C-D).**

**Figure 4:**
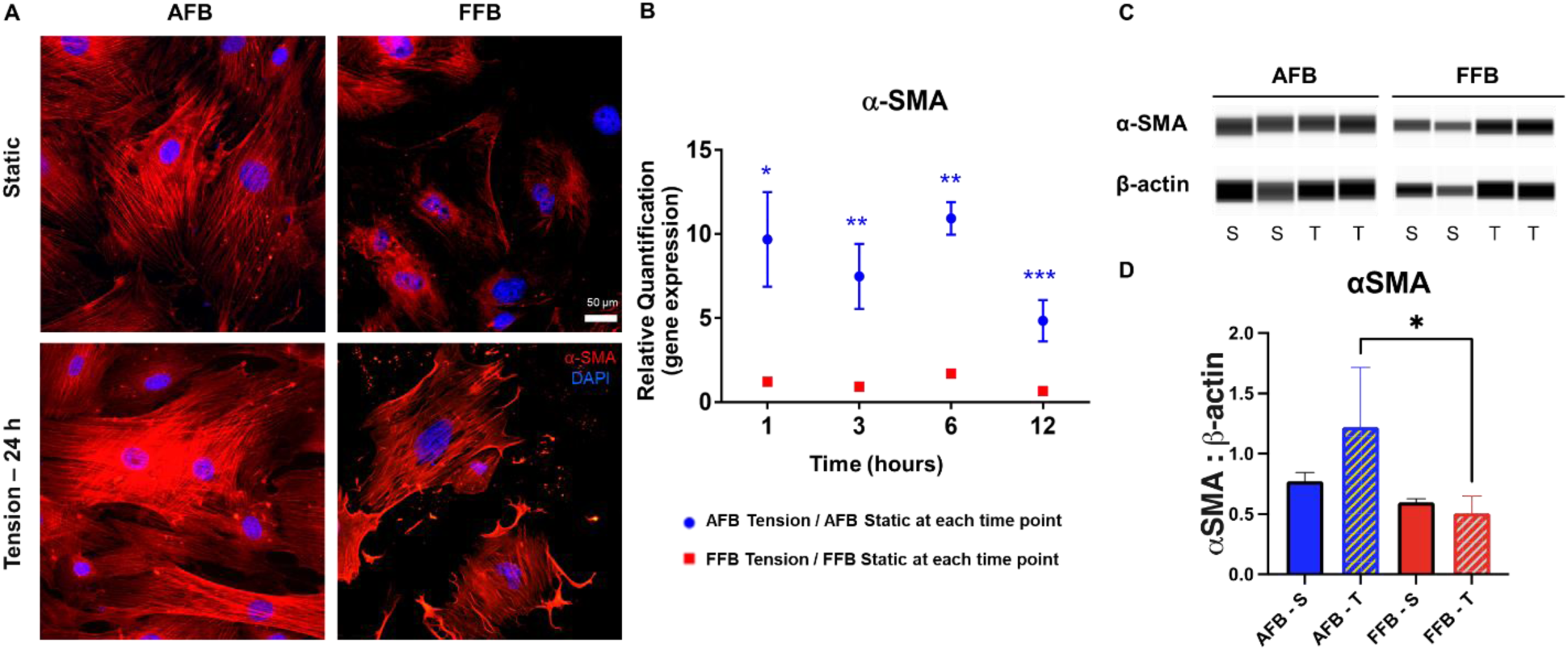
(A) Immunocytochemistry fluorescent staining of α-SMA (red) in AFB and FFB under static and 24 h tension conditions is shown. Blue stain indicates DAPI staining for cell nucleus. Scale bars represent 50 µm. n=3, with each experiment conducted with duplicate wells per group that were used one each for staining and staining control. Representative images from one experiment are shown. (B) Relative gene expression of α-SMA at 1, 3, 6, and 12 hours. Each time point is expressed as mean ± SD of gene expression of tension conditions normalized to paired static controls at that time point in each cell line. *p<=0.05, **p<=0.01, ***p<=0.001 denotes significant changes in tension compared to baseline expression in each cell line. Blue * indicates significance value for AFB, Red * indicates significance value for FFB. (C) Proteinsimple detection of α-SMA and β-actin from cell lysate of fibroblasts following static (S) or tension (T) conditions. (D) Quantification of α-SMA normalized to β-actin. N=3 experiments per treatment group.

### Tension differentially increases fibrogenic potential in both adult and fetal fibroblasts

Our collective data suggested that tension induces changes in HA expression and molecular weight, along with changes in the ECM organization and stress filaments, which could also lead to subsequent changes in the functional fibrotic phenotype of the FFB. To begin to understand how tension effects the fibrotic phenotype of FFB and AFB, we sought to evaluate CD26, a cell surface marker that has been shown to be positively correlated with fibroblast fibrogenic propensity ^2^. **Figure 5A** demonstrates representative immunofluorescent staining for CD26, which shows the AFB have more CD26 staining than FFB under static conditions. While there was an increase in CD26 expression in the FFB after 24 hours of tension than under static conditions, the CD26 expression of AFB appears to remain unchanged with tension. Analysis of CD26 gene expression revealed an upregulation in the FFB under tension at early time points at 1 and 3 h versus static conditions, with no significant change in expression over time in AFB. FFB had an almost 5-and 3-fold increase in CD26 expression under tension at 1 and 3 hours respectively (**Figure 5B**). These results suggest that 10% strain induced gene expression changes are consistent with a shift toward a fibrogenic phenotype in FFB.

**Figure 5:**
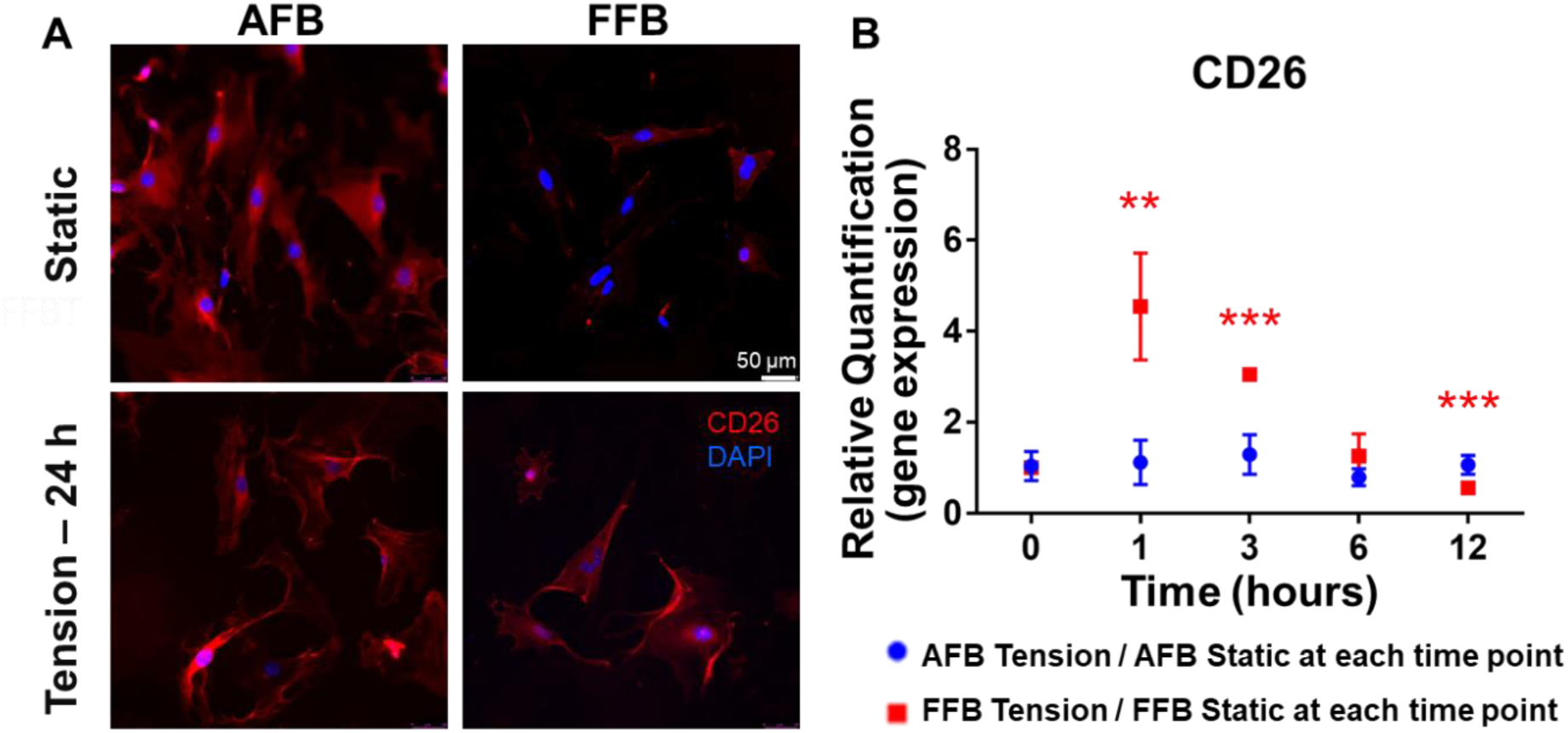
(A) Immunocytochemistry fluorescent staining of CD26 (red) in AFB and FFB under static and 24 h tension conditions. Blue stain indicates DAPI staining for cell nucleus. Scale bars represent 50 µm. n=3, with each experiment conducted with duplicate wells per group that were used one each for staining and staining control. Representative images from one experiment are shown. (B) Relative gene expression of CD26 at 1, 3, 6, and 12 hours. Each time point is expressed as mean ± SD of gene expression in tension conditions normalized to paired static controls at each time point in each cell line. *p<=0.05, **p<=0.01, ***p<=0.001 denotes significant changes in tension compared to baseline expression in each cell line. Blue * indicates significance value for AFB, Red * indicates significance value for FFB. N=3 experiments per treatment group.

To further characterize the effect of tension on fibrotic gene changes in FFB and AFB, we next utilized a PCR-based 84 gene fibrosis array to analyze expression patterns in AFB and FFB under static culture and after 6 hours of tension. Principal Component Analysis (PCA) of the genes ranked by their effect on overall fibrotic phenotype are plotted in **Figure 6A**, which clearly indicates a separation in AFB and FFB in static and tension conditions. In multi-dimensional sampling analysis, FFB under tension were more similar in phenotype to AFB than static FFB, indicating an effect of tension on inducing a fibrotic phenotype in FFB **(Figure 6B). Figure 6C** shows Venn diagrams illustrating the proportion of common or unique differentially up-or down-regulated fibrotic genes seen between AFB and FFB exposed to tension. We found 17 genes to be upregulated with tension: 5 of them exclusively in FFB, 4 exclusively in AFB, and 8 upregulated in both FFB and AFB. We found 28 genes to be downregulated with tension: 13 of them exclusively in FFB, 8 exclusively in AFB, and 7 downregulated in both FFB and AFB. The specific up-or down-regulated genes illustrated in **Figure 6C** are listed in **Table 3** (upregulated genes) and **Table 4** (downregulated genes). Many genes involved in inflammation (Interleukins 1a, 5, and 13) and inflammatory signal transduction (C-jun) were upregulated by tension in AFB and FFB. Itga2 and Itgb3 are genes involved in cell adhesion and were downregulated by tension in both AFB and FFB. Some genes involved in the TGF-β family of inflammatory signal transduction (Smad6 and Thbs1) were downregulated by tension in both AFB and FFB. A gene involved in ECM remodeling (MMP13) was exclusively upregulated in FFB exposed to tension. Expression of integrin subunits alpha 2 and beta 3 were downregulated in both AFB and FFB in response to tension, and subunits alpha 1 and beta 8 only in FFB.

### FFB cell morphology and adherence is more affected by tension than AFB cells

We observed that both the cellular morphology and density of only FFB monolayers appeared to change over time under mechanical tension, with a significant decrease in cell density of adhered cells after 24 h of tension, thus we sought to objectively measure the absolute change in cell number in AFB and FFB after tension. Cells were subjected to tonic 10% strain for up to 24 h similarly as prior experiments, and the morphology and cell attachment were assessed **(Figure 7A-J).** Adult fibroblasts tolerated tension and appeared normal in cell morphology, with minimal change in cell density after 24 h of exposure to tension. In contrast, after 6 h of exposure to tension, some FFB appeared rounded with disrupted attachment to the flexible membrane culture plate, and by 24 h, the absolute number of FFB adhered to the culture plate had decreased. Quantification of both AFB and FFB confirmed these observations **(Figure 7K).** Results are shown as a percent change from the controlled seeding density at 0 h prior to application of tension in each cell type. For AFB, there was minimal percent change in cell density throughout the 24 h period, with a slight increase (from 100% density to 109%. In contrast, there was a significant decrease in FFB cell density, with a mean density of 100% at 0 h, 76% at 3 h, 47% at 6 h, 19% at 12 h, and 10% by 24 h (p<0.001).

**Figure 6:**
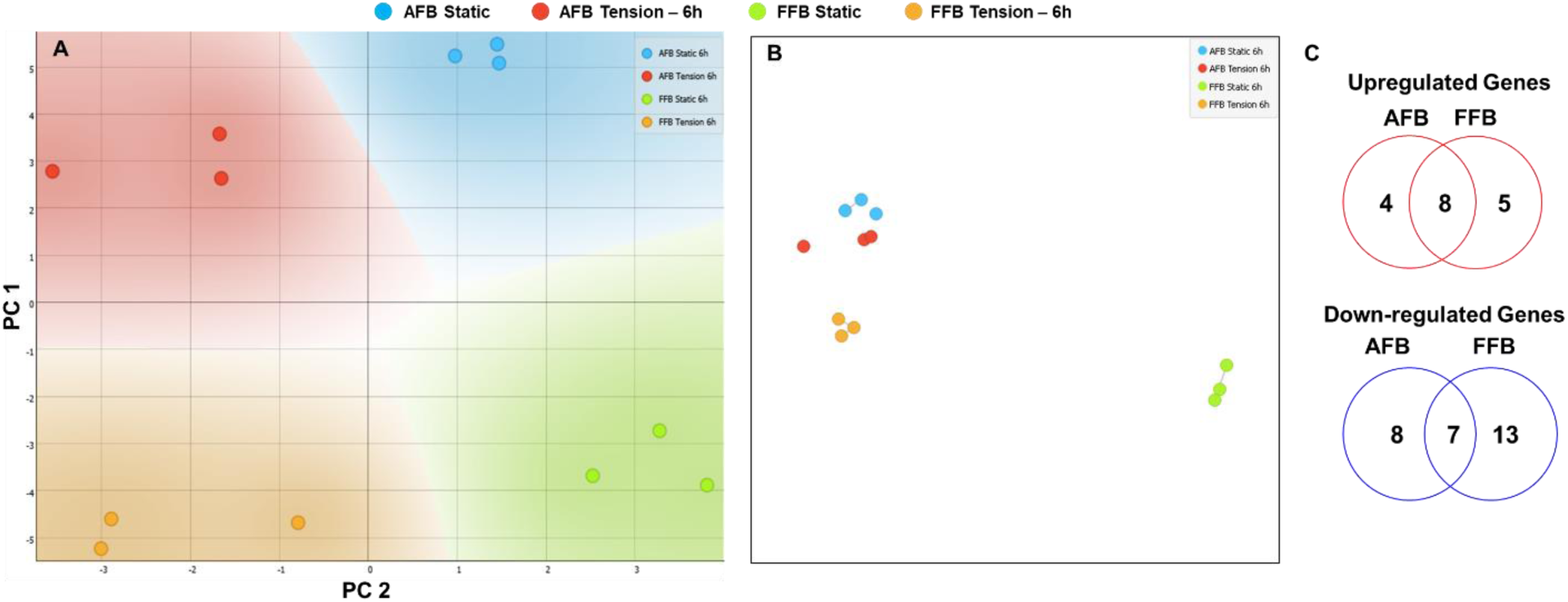
(A) Principal Component Analysis of the top 25 fibrosis genes that differentiate the AFB and FFB under static and 6 h tension conditions is shown. (B) Multi-dimensional scaling analysis of these genes shows a clear separation in the grouping of AFB and FFB under static culture conditions. (C) Venn diagrams indicate the total number of fibrotic genes that were up-or down-regulated in AFB and FFB cells.

**Figure 7:**
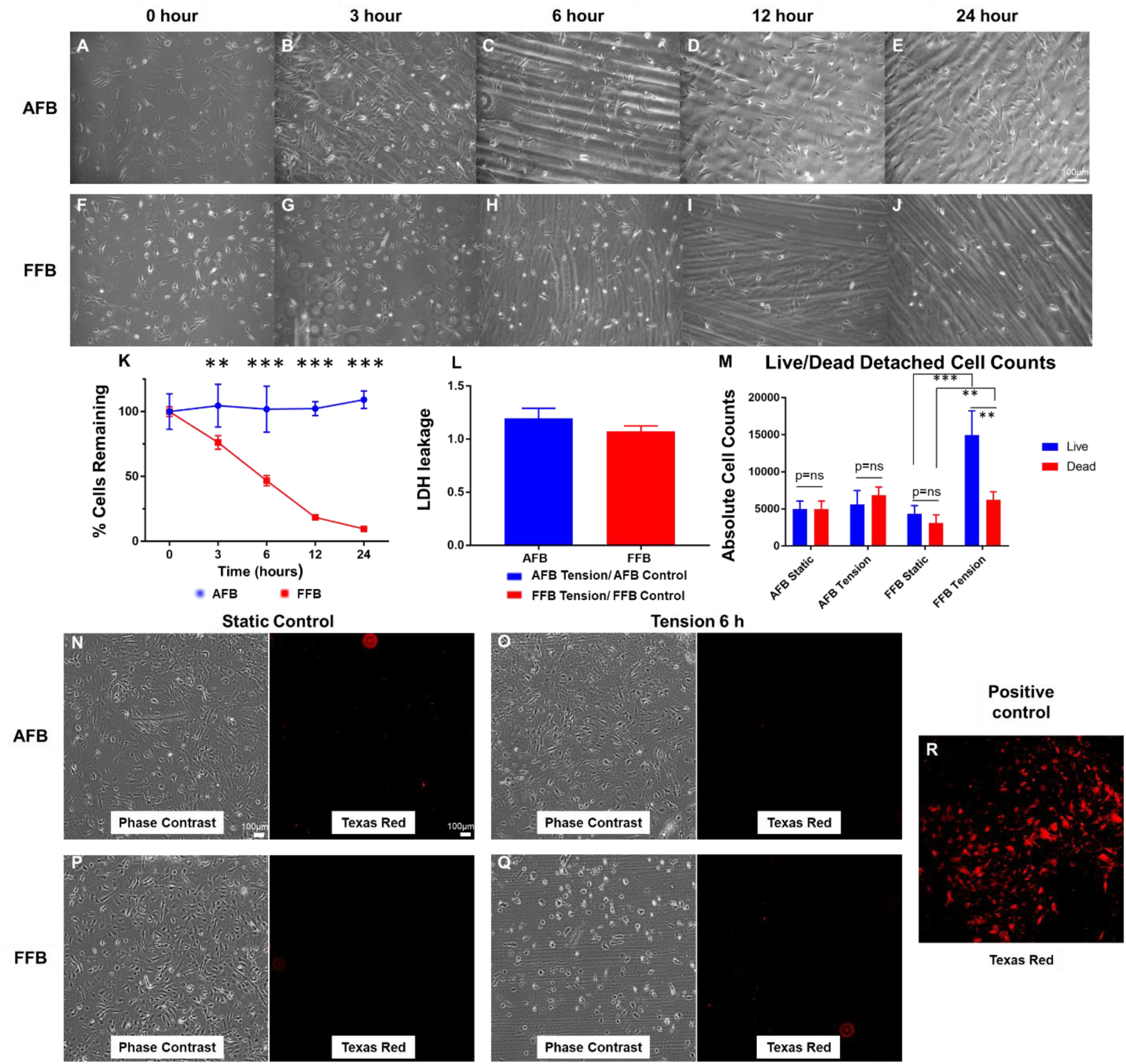
Phase-contrast images of AFB and FFB after 0 (A, F), 3 (B, G), 6 (C, H), 12 (D, I), and 24 (E, J) hours of 10% tonic tension. Streaks are due to vacuum grease residue that is utilized with the Flexcell machine. Scale bars represent 100µm. (K) Images were taken at 10X magnification and attached cells that had a well-spread, spindle shaped morphology were counted. Each timepoint is expressed as a % of cells remaining from T 0 ± SD. *p<=0.05, **p<=0.01, ***p<=0.001 denotes significant changes in % cells between AFB and FFB. (L) LDH levels in cell culture supernatant were evaluated after 6 h of static or 10% strain treatments. The levels in tension conditions were normalized to static culture for each cell type. Bar plot represents mean ± SD. (M) Quantification of Trypan blue cell counts. Supernatant was collected from wells after 6 hours of 10% tension and was stained with trypan blue and live/dead cells in the supernatant were counted and plotted as mean ± SD. *p<=0.05, **p<=0.01, ***p<=0.001 denotes significant changes. (N-Q) Immunocytochemistry fluorescent Live/Dead staining of AFB and FFB under static and 6 h 10% tension conditions. Phase contrast at 5X magnification (left) as well as fluorescent images of the same frame in the Texas red channel (right) are shown. (R) As a positive control, FFB subjected to excessive mechanical stress are shown that fluoresce highly in the Texas Red channel. The small red specks noted in the images is background and not localized to a cell. Similarly, the streaks in phase contrast images are remnants of vacuum grease captured in low magnification images that is used on the outer side of flexible culture membrane to reduce friction during the application of strain cycles, and does not indicate any membrane deformation. Scale bars represent 100 µm. N = 3 for each experimental condition.

In order to determine whether tension effected the viability of FFB cells (i.e. cell membranes were compromised and FFB were dying) or tension effected the adherence of the FFB (i.e. FFB were rounding up and detaching from the membrane under tension), we performed an LDH assay on the cell culture supernatants, along with counting live vs. dead cells floating in the supernatant with trypan blue uptake of dead cells. We also performed Live/Dead fluorescent staining of the remaining adherent cell monolayers on the flexible membranes. Results of the LDH assay are shown in **Figure 7L**. The LDH levels in the supernatant of cells subjected to 6 hours of 10% tension was normalized to the LDH levels in supernatant of static cells for each cell type. There were no significant differences in the expression between AFB and FFB (1.2 vs 1.1), suggesting that the cell membranes were probably not compromised by tension in either AFB or FFB at this time point, despite 50% of FFB no longer adhered **(Figure 7K).**

Live vs. dead cell counts in the cell culture supernatant based on trypan blue uptake **(Figure 7M)** revealed no difference in either live or dead cells in tension vs. static in AFB. We saw an increase in the total number of cells that were present in the supernatant of FFB cells after 6 h of tension vs. static control conditions. However, as compared to AFB, there were 3-fold more live cells in FFB after 6 h under tension, whereas the number of dead cells in FFB under tension was similar to that in AFB under tension. Lastly, we performed Live/Dead fluorescent staining **(Figure 7N-Q)**. There was no qualitative difference in the total fluorescence between AFB and FFB under either static or tension conditions, with no remarkable positive staining of cells in either group. However, FFB cells under tension appeared less dense and some showed a more rounded morphology in phase contrast imaging **(Figure 7Q).** Cells that had a disrupted membrane in positive control samples took up the stain and fluoresced more brightly than intact cells. Collectively, these data indicate that fetal cells are not dying after 6 hours of exposure to tension, but rather detaching from the membrane that is subjected to tonic tension.

## DISCUSSION

The wound repair process is influenced by many intrinsic and extrinsic factors. This study was aimed at investigating the differences in responses to mechanical tension between fetal and adult dermal fibroblasts, and to further understand the fetal regenerative vs. reparative healing phenotype marked by fibrosis in postnatal wounds. We specifically investigated the effect of tension on FFB, which naturally reside in a low mechanical stress environment, by measuring changes in hyaluronan metabolism, cell morphology and cytoskeletal components, and the markers of fibroblast fibrotic phenotype, as compared to the responses in AFB. Our results showed that FFB respond to tension by decreasing synthesis and increasing degradation of their predominant ECM component, HMW-HA, thus reducing their HA-rich PCM to resemble the PCM of the AFB counterparts. HA is a major structural and functional component of the extracellular matrix and plays an essential role in key FFB responses, such as their migration, wound healing, and invasion, that differentiate the FFB from AFB. It is posited that the hydrated microenvironment made possible by HA-rich ECM allows effective migration and invasion of the FFB during the regenerative wound healing phase, which is not prevalent in their postnatal counterparts. Instead, AFB utilize enzymatic cleavage to migrate through the granulation tissue in the wounds ^26^. The effect of tension on altering HA in FFB suggests that tension can alter the FFB cell phenotype to induce a more AFB-like fibrotic phenotype, which may affect their regenerative wound healing capability.

We also explored the balance of HA synthesis and degradation in AFB and FFB in static and tension conditions. To this end, we measured the expression levels of hyaluronan synthesis and degradation genes, and showed that FFB under tension have increases in LMW-HA production genes and degradation genes while AFB show decreased HMW-HA production and increased expression of hyaluronan degradation enzymes, both profiles indicating an overall shift towards a LMW-HA profile under tension. While our finding of a decrease in TMEM2 expression, known to cleave HA into medium sized fragments, with tension does not fit the profile of a shift towards LMW-HA, recent findings suggests that TMEM2 does not act as a functional hyaluronidase in skin fibroblasts ^31, 32^. The same group has demonstrated that knockdown of TMEM2 enhanced HA degradation and increased expression of KIAA1199 in human skin fibroblasts ^33^. Therefore, based on its inverse relationship with KIAA1199, the decrease of TMEM2 expression may contribute to the degradation of HMW-HA seen here by upregulating KIAA1199 expression in our FFB. Certain cytokines such as IL-6 and IL-8, which are increased in larger fetal dermal wounds, have recently been shown to increase HA turnover via increased expression of KIAA1199, thus our results suggest that changes in HA metabolism in large fetal wounds with more tension may not only be a consequence of a pro-inflammatory cytokine milieu, but also the intrinsic effects of tension on FFB [33473125]^31^. Our data confirms previous reports of increased hyaluronidase activity as a characteristic of adult and late gestational fibrotic wound healing ^34^. This change in the homeostatic balance of HA turnover also corroborates our findings of overall decreased HMW-HA expression by FFB in response to tension. It has been proposed that prolonged presence of HMW-HA in fetal wounds may provide the mesenchymal signal for healing by regeneration ^35^; therefore, its loss may signal the formation of fibrosis and scar, as seen in critically large fetal wounds, where increased wound size causes increased mechanical forces that have to be overcome to close the wound.

Another important finding of this study is that the FFB, which were able to tolerate tension conditions expressed CD26, a marker of a profibrotic engrailed-1 expressing lineage of fetal fibroblasts ^3^. Mascharak et al. also recently demonstrated that mechanical tension on wounds leads to an increase in fibroblast differentiation into this engrailed-1 positive pro-fibrotic lineage ^17^. Our data similarly demonstrates, *in vitro,* that CD26 expression was upregulated in FFB only under tension, while AFB have elevated CD26 levels at baseline under static culture conditions, which was not affected by tension conditions. This upregulation of CD26 in FFB under tension suggests that only the profibrotic engrailed-1 FFB lineage is able to tolerate increases in microenvironmental tension, and thus remain attached to the culture membrane. While it is known that increasing wound size causes fetal scar formation via amplification of the inflammatory response through recruitment of inflammatory cells and expression of proinflammatory cytokines ^3, 4^, changes in microenvironmental tension in these large wounds may be another important factor in selectively promoting CD26 positive fibroblast activity and scar formation. Even though engrailed positive FFB could tolerate this stressor, engrailed-negative FFB may not function under these conditions of increased tension, which needs to be further explored. Furthermore, during development, the increase in CD26 expression coincides with a shift in wound healing from fetal regenerative to a reparative phenotype typical of postnatal and adult wounds. This also corresponds with a shift in the HA content of the wound ECM, as we have shown previously ^19^, and an increase in the resting tension of skin ^36^. Interestingly,

SMAD-6 was downregulated in both AFB and FFB under tension. SMAD-6 acts to disrupt the TGF-β downstream signaling pathway by preventing phosphorylation of SMAD-2, a canonical pathway through which TGF-β exerts fibrotic effects ^37^. Fetal skin is known to have a higher ratio of TGF-β3 to TGF-β1, physiology which contributes to the regenerative healing phenotype seen in fetal wounds ^38–40^. Downregulated SMAD-6, combined with the observed downregulation of anti-fibrotic TGF-β3 expression in FFB supports that tension induces a pro-fibrotic state in these fibroblasts. Notably, expression of four integrins were captured as being downregulated in our fibrosis array: subunits alpha 2 and beta 3 were downregulated in both AFB and FFB in response to tension, and subunits alpha 1 and beta 8 only in FFB. Integrins with an alpha 1 or 2 subunit play a large role in collagen binding, downregulation of which may have led to the cellular detachment from the membranes seen by FFB ^41^. In particular, binding of alpha 1 beta 1 integrins to collagen downregulates collagen gene expression, while alpha 1 beta 2 integrin binding can upregulate matrix metalloproteinase expression ^42^. Tension’s effects on integrin expression in FFB may alter the turnover and organization of ECM and lead to the increased fibrosis seen in larger fetal wounds. Notably the expression in collagen genes did not change markedly in either FFB or AFB. Though many studies do use changes in collagen synthesis as a measure of fibrotic ECM production, recent studies posit that the quantity of collagen produced in a wound does not alter the scarring outcome so much as the specific orientation of the collagen fibrils ^17^. While previous studies suggest that mechanical tension, ECM HA content, and fibroblast phenotype are intertwined in producing putative regenerative vs. reparative responses, our study provides a clearer understanding of the regulation of the FFB ECM and fibrotic phenotype by tension.

We also assessed the levels of α-SMA, a marker of myofibroblasts, characteristic of adult wound healing. α-SMA is negligible in fetal wound healing, and its appearance coincides with the onset of scar formation seen in later gestation ^43^. While tension had no effect on α-SMA gene expression or protein levels in FFB, qualitative analysis of α-SMA organization demonstrated that the intracellular α-SMA in FFB under tension resulted in parallel cords that spanned the entire cell, resembling the structure in AFB. The same observations of α-SMA organizational structure were seen with F-actin staining, which changed from shorter disorganized fibers at baseline in FFB to parallel bundles that span the entire cell under tension, akin to that in the AFB. These organized bundles represent stress fibers, which consist of numerous actin filaments, and can produce contractile forces similar to myofibrils of muscle cells, though lacking the same organization ^44^. While chemical signaling pathways are known to regulate the formation of stress fibers, the ultrastructure of these fibers are dynamic, and their formation and orientation can be influenced by intra-and extracellular mechanical forces ^45^. The observation that tension induces a change in the organization of stress fibers in FFB does not necessarily signal a change to a more fibrotic phenotype; however, it is known that the function of stress fibers are, in part, determined by their ultrastructure, and range from contraction to transcriptional regulation of genes involved in matrix reorganization and cell locomotion ^46^. Indeed, we noticed that FFB under tension appear to change their morphology from a spread spindle shape to a more rounded cell morphology. Furthermore, a majority of these cells lift off from the membranes. Since our live/dead analysis of cells still attached on the membranes and the LDH analysis and cells counts in the supernatant indicate no cell membrane disruption, but rather more floating live cells in FFB under tension, one possible explanation is that these cells cannot adhere under tension. Our gap in study of these free-floating FFB presents a limitation of this work; exploring the phenotypic and genomic differences of these cells unable to tolerate tension is an area of further work we are pursuing. The concomitant changes in cytoskeletal organization and HA cable-like structures in FFB under tension suggest that these structural changes regulate how these cells sense and respond to the microenvironment. Though it is beyond the scope of this paper to characterize the ultrastructure and function of the stress fibers in governing the FFB phenotype, this is a future testable hypothesis.

Our finding that tension reduces the number of FFB adherent to the culture plate poses a limitation on the interpretation of our data. We see an increase in CD26 expression in FFB under tension, however, the experimental design does not allow us to determine if only the CD26 positive lineage is able to tolerate tension, or if tension is inducing CD26 in all cells and some of those cells happen to detach from the culture plate. In the context of fetal wound healing, knowing that CD26 is a marker of a pro-fibrotic fibroblast lineage, it stands to reason that the mechanical force generated within a larger wound in the fetus may create an environment only CD26 positive fibroblasts can tolerate. Thus, the scarring seen in large fetal dermal wounds may be attributed to a lineage of fibroblasts that both tolerates tension and has a fibrotic phenotype, while the lineages of non-CD26 positive anti-fibrotic fetal fibroblasts are unable to tolerate the increased mechanical strain in the larger wound environment or contribute to its repair.

A closer look at the time course of CD26 expression in FFB under tension reveals that CD26 is most upregulated at 1 and 3 hours of tension, relatively early in the time course of the experiment. At these time points, there has not been a dramatic decrease in FFB cell numbers. We posit that tension does trigger increased expression of CD26, but only those cells that are further down the path of differentiation into CD26 positive cells are able to confer the fibrotic phenotype that allows for tolerance of the tension. Further mechanistic studies are necessary to address this dilemma.

In conclusion, our data demonstrate a differential response to tension between fetal and adult fibroblasts when exposed to increased tension. Namely, tension alone is able to shift the regenerative, HMW-HA producing phenotype of fetal fibroblasts towards an adult-like pro-fibrotic phenotype, which could explain why scarring is seen in fetal wounds of greater size. Scarring is a disease process that causes significant morbidity and mortality, and though currently available anti-scarring therapies make up a multi-billion-dollar industry, none have been able to successfully reverse the scarring phenotype ^47, 48^. Understanding the mechanisms behind the fetal regenerative phenotype, specifically those that permit regeneration and those that hinder the regenerative capabilities will allow us to pursue anti-scarring therapeutics based upon the elucidation of fundamental biological processes.

## INNOVATION

Recent works have isolated mechanotransduction pathways as crucial to activating the fibrotic response in wound healing that leads to scarring. However, there remains a gap in knowledge with regards to how tension affects fetal wound healing, thus reversing the fetal regenerative phenotype. Our data begin to investigate tension in fetal wound healing in novel ways by examining its effects on fetal fibroblasts, particularly with regards to hyaluronan metabolism. These and further studies will continue to unravel the precise underpinnings of fetal wound healing, thereby allowing for the development of novel wound healing therapeutics.

## KEY FINDINGS

- Static tension alters the profile of hyaluronan metabolism of FFB, shifting them towards reduced production of HMW-HA and degradation of HA by hyaluronidases into LMW-HA.
- FFB grown in a high-tension environment have less pericellular matrix, similar to AFB, which may impact their functional phenotype.
- Tension differentially increases the fibrogenic potential in both adult and fetal fibroblasts.
- FFB membrane adherence is impaired by tension, however this did not result in fibroblast cell death.

## ACKNOWLEDGMENTS AND FUNDING SOURCES

We would like to thank Dr. Numan Oezguen, Department of Pathology, Texas Children’s Hospital and Baylor College of Medicine for training in Orange Software. We would also like to acknowledge Roma Patel and Swetha Saravanan for their help with imaging and analyzing data. We thank Dr. Jennifer McCracken, and Dr. Hector Martinez-Valdez in the Office of Surgical Research Administration in the Department of Pediatric Surgery, Texas Children’s Hospital for manuscript editing. This research was supported by the John S. Dunn Foundation Collaborative Research Award Program administered by the Gulf Coast Consortia, 3M Award given by the Wound Healing Foundation, Clayton Seed Grant by the Department of Pediatric Surgery, Texas Children’s Hospital, and the Interim funding by Baylor College of Medicine, and NIGMS-R01GM111808.

## AUTHOR DISCLOSURE AND GHOSTWRITING

SB, MMF, WDS, UMP and NT designed the experiments. UMP, NT, AB, DC, HS, AK, HVV, HL, MMF, OOO, PB, PK, OSJ and SB performed cell isolation and bench experiments. UMP, WDS, NT, MMF, BP, CC, OOO, PB, PK, OSJ and SB analyzed data. UMP, WDS, BP, MF, OOO, and SB wrote the manuscript. SB, CC, and MMF provided scientific comments for the study and significant content revisions to the manuscript. All authors have reviewed and approve the final version of the manuscript. The authors declare that they have no competing interests. This article was written by the authors and ghostwriting services were not used.

## ABOUT THE AUTHORS

**Walker Short, MD** is a surgical resident at Baylor College of Medicine (BCM) who completed a 2-year Postdoctoral Research Fellowship in the Laboratory of Regenerative Tissue Repair (LRTR), who focuses on wound healing, inflammation, and fibrosis. **Umang Parikh** and **Daniel Colchado** are former medical students who completed predoctoral fellowships with the LRTR. **Natalie Templeman**, **BS**, **Alexander Blum, BS, Benjamin Padon, BA** and **Aditya Kaul, BS** are former predoctoral fellows in the LRTR. **Oluyinka O. Olutoye II, MD, MPH** is a surgical resident at Baylor College of Medicine (BCM) completing a 2-year Postdoctoral Research Fellowship in the (LRTR), who focuses on wound healing and neonatal pulmonary hypertension. **Pranav Bommekal** and **Philip Kogan** and undergraduate interns in the LRTR, and **Olivia S. Jung** is a high school student who conducted summer research in the LRTR. **Hui Li, PhD** is a research associate in the LRTR. **Hima Vangapandu, PhD** and **Monica Fahrenholtz, PhD** are former Postdoctoral Fellows in the LRTR. **Cristian Coarfa, PhD** is an Associate Professor in the Department of Molecular and Cellular Biology at BCM. **Swathi Balaji, PhD** is an Assistant Professor in the Department of Surgery at BCM.

### LIST OF ABBREVIATIONS

HA: Hyaluronan
EN1: Engrailed-1
ECM: Extracellular Matrix
MW: Molecular Weight
LMW-HA: Low Molecular Weight Hyaluronan
HMW-HA: High Molecular Weight Hyaluronan
HAS: Hyaluronan Synthase
HYAL: Hyaluronidase
FFB: Fetal Fibroblasts
AFB: Adult Fibroblasts

## Supplemental Figure 1

**Supplemental Figure 1:**
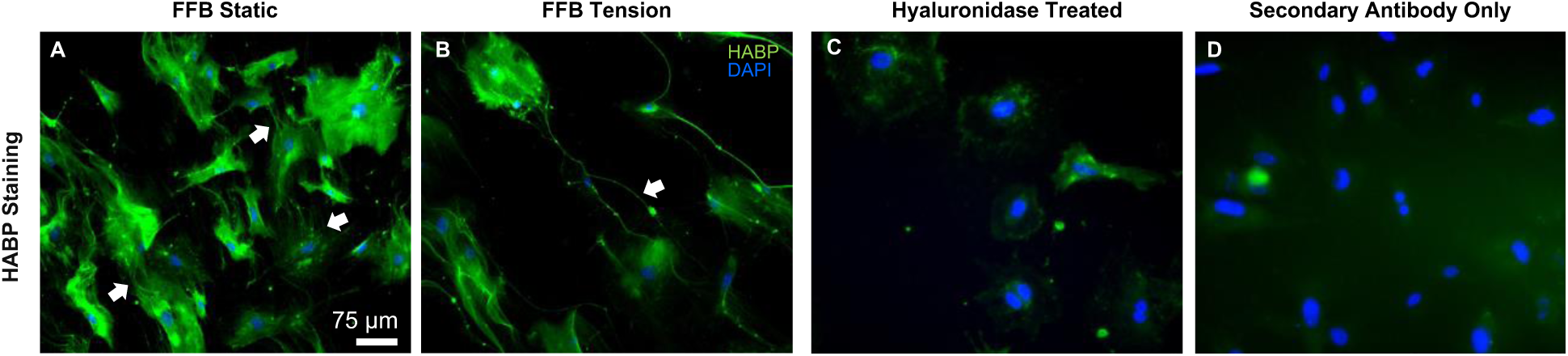
(A-B) Immunocytochemistry fluorescent staining of HA with HA binding protein (HABP, green) and DAPI for cell nucleus (blue). In the FFB under static conditions, HA is diffusely expressed within the cell, and there were distinct HA cable-like protrusions forming on each cell that connected to the neighboring cells (indicated by white arrows). After 6 h of tension, these cells appear to have down-regulated HABP expression intensity as compared to static conditions, and the HA cables appeared to form thick, cord-like strands between cells (indicated by white arrows). (C-D) Hyaluronidase-treated and secondary-only immunofluorecent images for specificity. Scale bars represent 75 µm.

