## supplemental figure for "Phenotypic Differences in Adult and Fetal Dermal Fibroblast Responses to Mechanical Tension"

**Supplemental Figure 1**

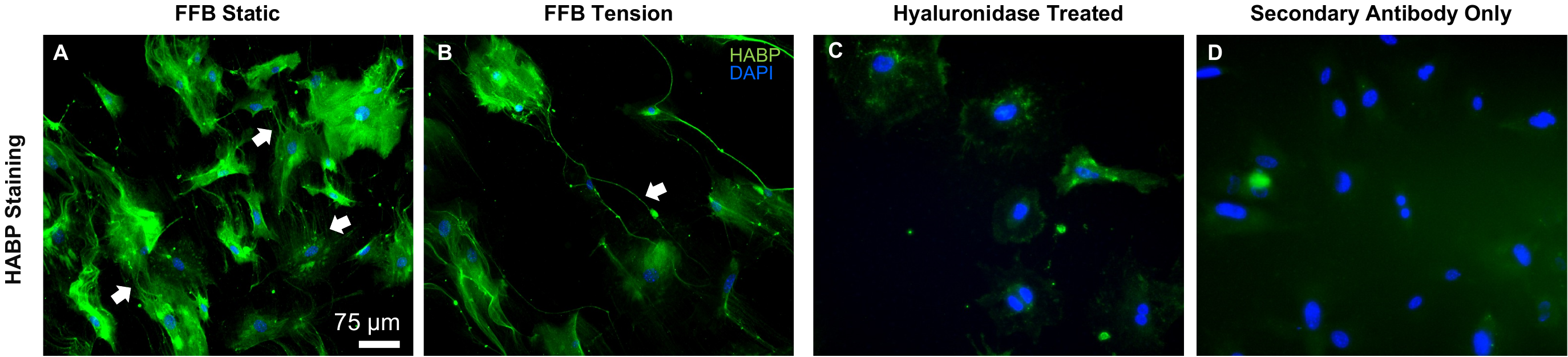

**Supplemental Figure 1:** (A-B) Immunocytochemistry fluorescent staining of HA with HA binding protein (HABP, green) and DAPI for cell nucleus (blue). In the FFB under static conditions, HA is diffusely expressed within the cell, and there were distinct HA cable-like protrusions forming on each cell that connected to the neighboring cells (indicated by white arrows). After 6 h of tension, these cells appear to have down-regulated HABP expression intensity as compared to static conditions, and the HA cables appeared to form thick, cord-like strands between cells (indicated by white arrows). (C-D) Hyaluronidase-treated and secondary-only immunofluorecent images for specificity. Scale bars represent 75 µm.
